# The zinc finger antiviral protein ZAP destabilises viral transcripts and restricts human cytomegalovirus

**DOI:** 10.1101/2020.09.15.297804

**Authors:** Ana Cristina Gonzalez-Perez, Markus Stempel, Emanuel Wyler, Christian Urban, Antonio Piras, Thomas Hennig, Albert Heim, Markus Landthaler, Andreas Pichlmair, Florian Erhard, Lars Dölken, Melanie M. Brinkmann

**Affiliations:** Viral Immune Modulation Research Group, Helmholtz Centre for Infection Research, Inhoffenstr. 7, 38124 Braunschweig, Germany; Institute of Genetics, Technische Universität Braunschweig, Spielmannstr. 7, 38106, Braunschweig, Germany; Berlin Institute for Medical Systems Biology, Max-Delbrück-Center for Molecular Medicine, 13125 Berlin, Germany; School of Medicine, Institute of Virology, Technical University of Munich, Schneckenburgerstr. 8, 81675 Munich, Germany; Institute for Virology and Immunobiology, Julius-Maximilians-Universität Würzburg, Versbacher Str. 7, 97078 Würzburg, Germany; Helmholtz-Institute for RNA-based Infection Research, HIRI, Josef-Schneider-Straße 2/D15, 97080 Würzburg, Germany; Institute of Virology, Hannover Medical School, Carl-Neuberg Str. 1, 30625 Hannover, Germany; IRI Life Sciences, Institute of Biology, Humboldt-Universität Berlin, Philippstraße 13, 10115 Berlin, Germany

**Keywords:** ZAP, ZC3HAV1, ISG, antiviral, DNA virus, herpesvirus, HCMV, innate immunity

## Abstract

Interferon-stimulated gene products (ISGs) play a crucial role in early infection control. The ISG zinc finger CCCH-type antiviral protein 1 (ZAP/ZC3HAV1) antagonises several RNA viruses by binding to CG-rich RNA sequences, whereas its effect on DNA viruses is largely unknown. Here, we decipher the role of ZAP in the context of human cytomegalovirus (HCMV) infection, a β-herpesvirus that is associated with high morbidity in immunosuppressed individuals and newborns. We show that expression of the two major isoforms of ZAP, the long (ZAP-L) and short (ZAP-S), is induced during HCMV infection and that both negatively affect HCMV replication. Transcriptome and proteome analyses demonstrated that the expression of ZAP decelerates the progression of HCMV infection. SLAM-sequencing revealed that ZAP restricts HCMV at early stages of infection by destabilising a distinct subset of viral transcripts with low CG content. In summary, this report provides evidence of an important antiviral role for ZAP in host defense against HCMV infection and highlights its differentiated function during DNA virus infection.

## Introduction

Viral infections pose a major global health burden as the cause of a range of debilitating human diseases with the potential to paralyse countries. Herpesviruses are large, structurally complex DNA viruses belonging to the *Herpesviridae*. Within this family, a number of viruses are responsible for a variety of diseases in humans ranging from cold sores and pneumonia to cancer. The common peculiarity of herpesviruses lies in their ability to establish latency, which presents a great challenge in medicine due to severe complications resulting from virus reactivation. Human cytomegalovirus (HCMV) is one of the nine human herpesviruses described to date and the prototype virus of the *Betaherpesvirinae* subfamily. HCMV displays a coding capacity that far exceeds that of most other *Herpesviridae*, having the largest genome among all known human viruses and the capacity to encode more than 200 proteins. Primary HCMV infection generally causes mild symptoms in immunocompetent individuals (Cohen & Corey, 1985). However, immunosuppressed individuals, such as AIDS patients or transplant recipients, are vulnerable to HCMV-related disease (Meyers *et al*, 1986; Zamora, 2004). In addition, HCMV is the leading cause of congenital viral infection worldwide, and can result in serious long-term sequelae in newborns such as hearing loss, vision abnormalities, microcephaly, or developmental delays (Ramsay *et al*, 1991).

The host innate immune system, as the first line of defence, is equipped with germline-encoded pattern recognition receptors (PRRs), a group of sensors that detect pathogens by recognising pathogen-associated molecular patterns (PAMPs). The detection of PAMPs induces downstream signalling culminating in the activation of several transcription factors including interferon regulatory factors (IRF) and nuclear factor kappa-light-chain-enhancer of activated B cells (NF-κB), leading to the induction of genes encoding for type I interferons (IFNs), proinflammatory cytokines and non-canonical interferon-stimulated genes (ISGs) (Schoggins *et al*, 2014). Upon binding type I IFNs, the interferon-α/β receptor (IFNAR) is activated and its signalling results in nuclear translocation of STAT1 and STAT2 transcription factors and induction of canonical ISGs (reviewed in Schneider *et al*, 2014). ISGs are essential antiviral effectors and constitute a group of cellular factors ranging from PRR (e.g. IFI16, cGAS or RIG-I) or transcription factors to pro-apoptotic proteins or proteins involved in the regulation of the immune response (Gonzalez-Perez *et al*, 2020).

The zinc finger CCCH-type antiviral protein 1, also known as ZAP, ZC3HAV1 or PARP13, belongs to the subset of non-canonical ISGs whose expression can be induced via IRF3 directly as well as canonically by IFNAR signalling (Schoggins *et al*., 2014). Four isoforms of ZAP that originate from alternative splicing of the *ZC3HAV1* gene have been reported thus far (Li *et al*, 2019), with the long (ZAP-L) and the short (ZAP-S) isoforms being the most prominent ones. While approximately 700 amino acids are shared by ZAP-L and ZAP-S, ZAP-L has an extended C-terminus of around 200 amino acids containing a catalytically inactive PARP-like domain (Kerns *et al*, 2008) and a functional CaaX prenylation motif (Charron *et al*, 2013). The farnesyl modification on the cysteine residues of the CaaX motif increases the hydrophobicity of ZAP-L, targeting this isoform to membranes in endolysosomes (Charron *et al*., 2013). Both ZAP isoforms are equipped with an N-terminal zinc finger domain (containing four CCCH-type zinc finger motifs), a TiPARP homology domain (TPH), which is well conserved among ZAP paralogs and contains a fifth zinc finger motif, and a WWE domain, predicted to mediate specific protein-protein interactions (Aravind, 2001; Katoh & Katoh, 2003).

ZAP exhibits broad antiviral activity against a variety of RNA viruses by binding RNA and mediating its degradation (Guo *et al*, 2007). The antiviral activity of ZAP was demonstrated against alphaviruses (Bick *et al*, 2003), filoviruses (Muller *et al*, 2007), retroviruses (Takata *et al*, 2017; Zhu *et al*, 2011), and flaviviruses (Chiu *et al*, 2018). However, ZAP fails to inhibit a diverse range of other RNA viruses including vesicular stomatitis virus (VSV), poliovirus (Bick *et al*., 2003), influenza A virus (Liu *et al*, 2015; Tang *et al*, 2017), or enterovirus A71 (Xie *et al*, 2018). The involvement of ZAP in the defence against DNA viruses has not been explored to the same extent as for RNA viruses. While ectopic expression of ZAP failed to inhibit growth of the α-herpesvirus herpes simplex virus type 1 (HSV-1) (Bick *et al*., 2003), ZAP could restrict HCMV by an unknown mechanism (Lin *et al*, 2020). Interestingly, a luciferase-based reporter assay identified the HSV-1 UL41 protein, known for its ability to mediate degradation of several mRNAs, as a ZAP antagonist that degrades *ZAP* mRNA, which may explain why ZAP cannot restrict HSV-1 (Bick *et al*., 2003; Su *et al*, 2015). Modified vaccinia virus Ankara (MVA) was recently shown to be restricted by ZAP, and while the knockout of ZAP had no discernible effect on viral DNA, individual mRNA or protein species, an interference of ZAP with a late step in the assembly of infectious MVA virions was suggested (Peng *et al*, 2020).

To date, the RNA motif that is recognised by ZAP is still controversial. Early publications suggest an RNA structure-dependent recognition, based on the RNA tertiary structure, but already advocating the importance of the sequence-specific interaction between ZAP and its target RNA (Huang *et al*, 2010). The formation of tertiary structures raises the possibility of multiple binding sites. Subsequent studies support the recognition of CG-rich dinucleotide regions (Takata *et al*., 2017). Indeed, a recent study revealed on a structural level that ZAP binds to CG-rich RNA with high affinity through its basic second zinc finger, which contains a pocket capable of accommodating CG-dinucleotide bases (Meagher *et al*, 2019). However, another possibility is the binding of ZAP to AU-dinucleotides. Although one study claimed that ZAP does not recognise any of the three types of AU-rich elements (AREs) and concluded that ZAP may modulate stability of non-ARE-containing mRNAs (Guo *et al*, 2004), recent publications described ZAP target specificity for sequences enriched for AU-rich dinucleotides (Odon *et al*, 2019; Schwerk *et al*, 2019). Taken together, while there is solid data indicating ZAP recognition of CG-dinucleotide containing RNA, it is feasible that the presence of several other zinc finger motifs in the ZAP protein broadens its target specificity. How these target specificities are regulated and the mechanism of how ZAP-bound RNA is degraded warrants further investigation.

Here, we report that the expression of the ZAP-S and ZAP-L isoforms is induced upon infection and that both act as restriction factors for HCMV in human primary fibroblasts (HFF-1). Further, a combination of transcriptomics and proteomics demonstrates that ZAP restricts the progression of HCMV gene expression. Employing metabolic RNA labelling (SLAM-sequencing), we show that ZAP specifically affects the stability of distinct viral transcripts with low CG-content, namely transcripts of the RL11 gene family and the *UL144* gene, that are expressed with immediate-early to early kinetics of the viral life cycle. Altogether, we provide evidence that ZAP is an important cellular factor that restricts HCMV by a distinct manner.

## Results

### Expression of ZAP is induced upon HCMV infection

A previous proteomics study showed that HCMV infection of human fibroblasts (HFF-1) leads to the upregulation of a specific set of host proteins in the first 24 hours, including some ISGs. One of these ISGs was the antiviral protein ZAP (Weekes *et al*, 2014). The two major isoforms of ZAP, the short (ZAP-S) and the long (ZAP-L), are only differentiated by the inclusion of the C-terminal PARP domain in ZAP-L (Figure 1A). To analyse which of the two major isoforms of ZAP is induced upon HCMV infection, HFF-1 cells were infected with HCMV or, as control, stimulated with recombinant IFNβ, and expression of ZAP was analysed by immunoblotting. Our results show that ZAP-L was expressed in uninfected HFF-1 cells, and its expression was only marginally increased upon HCMV infection or IFNβ 24 hours post stimulation (Figure 1B). In contrast, ZAP-S was barely expressed in untreated cells, but its expression was strongly induced by HCMV infection and IFNβ treatment (Figure 1B). These results show that ZAP-S is a prototypical ISG, which is strongly induced upon HCMV infection, whereas ZAP-L is already expressed prior to, and only slightly induced with, infection.

**Figure 1.**
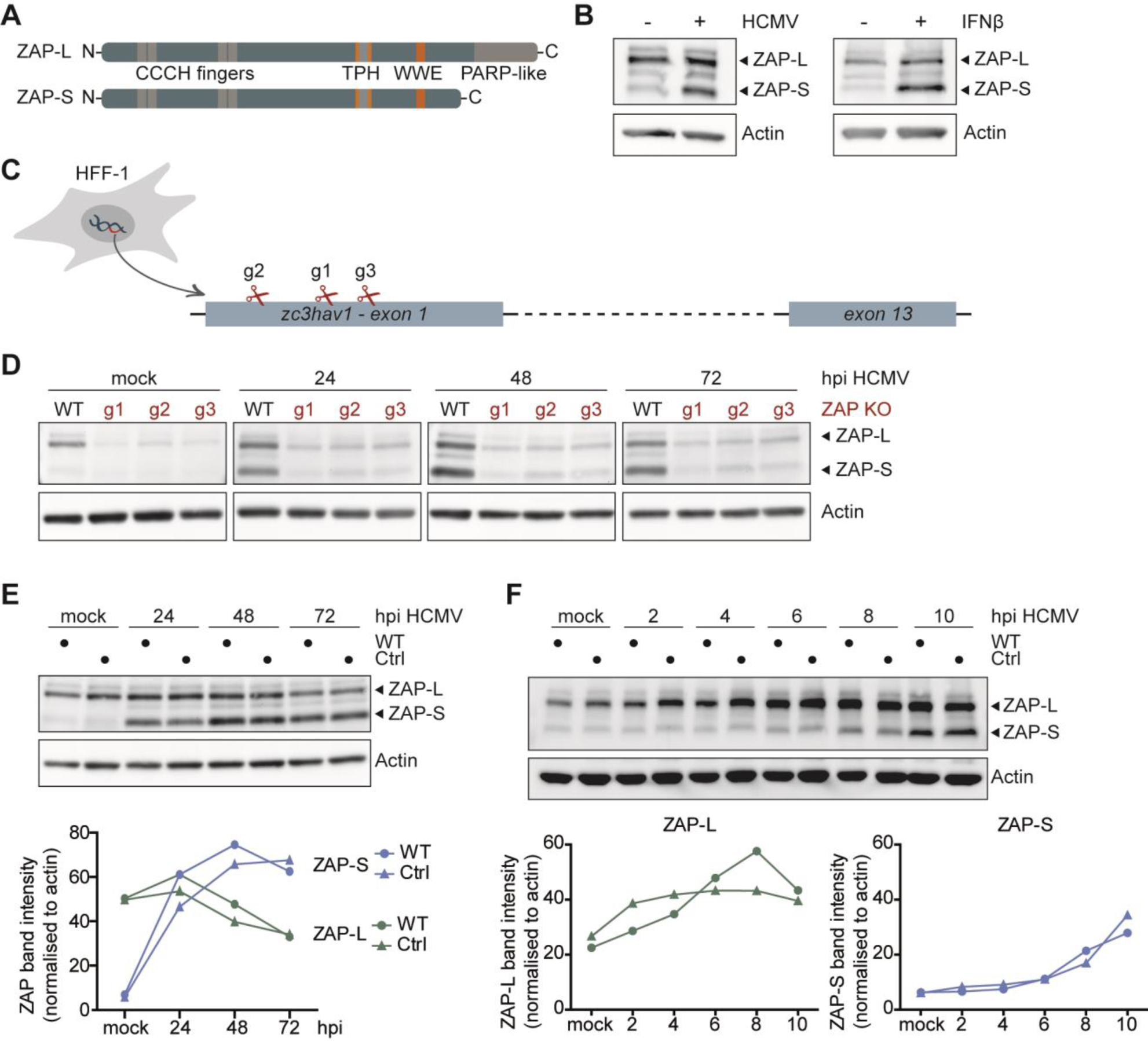
Expression kinetics of ZAP-S and ZAP-L in HCMV infected fibroblasts. **(A)** Schematic representation of the protein domains of the two main isoforms of ZAP, the long isoform ZAP-L and the short isoform ZAP-S. Both isoforms share four CCCH-type zinc finger motifs at the N-terminal domain, as well as a TPH domain containing a fifth zinc finger motif, and a WWE domain, while the C-terminal PARP-like domain is only present in ZAP-L. TPH = TiPARP homology domain, PARP = poly(ADP-ribose)-polymerase. **(B)** Primary human fibroblasts (HFF-1) were either mock treated, infected by centrifugal enhancement with HCMV (MOI 0.1), or stimulated by addition of recombinant IFNβ (20 ng/ml), and expression of ZAP and actin was analysed 24 hours later by immunoblotting with a ZAP- or actin-specific antibody. Hpi = hours post infection. **(C)** Three independent ZAP KO cell lines were generated by Cas9-mediated gene editing using three different gRNAs (g1, g2 and g3) which target the first exon of the *zc3hav1* gene. **(D-F)** Wild-type HFF-1 and the three ZAP KO **(D)** or control **(E-F)** cell lines were mock treated or infected by centrifugal enhancement with HCMV (MOI 0.1) for the indicated time points and cell lysates were subjected to immunoblotting with specific antibodies against ZAP and actin. Quantifications of ZAP-L (in green) or ZAP-S (in blue) band intensities normalised to actin are represented in line graphs.

To investigate whether ZAP can shape the course of HCMV infection, we generated three individual ZAP-deficient HFF-1 cell lines by Cas9-mediated gene editing. Each cell line was generated using a different guide RNA, all targeting the first exon of *Zc3hav1*, and thereby affecting both ZAP-L and ZAP-S expression (Figure 1C). To verify the efficacy of genome editing, we infected the knockout cell lines with HCMV and analysed ZAP expression at 24, 48, and 72 hours post infection (hpi). Protein levels of both ZAP-L and ZAP-S were strongly reduced in all three knockout cell lines (g1, g2, g3), confirming successful genome editing (Figure 1D). As a control, we also generated a cell line with a non-targeting guide RNA. Expression of ZAP followed the same kinetics in both wild-type (WT) and control HFF-1 cells, demonstrating that stable expression of Cas9 did not affect ZAP expression kinetics (Figure 1E). To pinpoint when ZAP-S expression begins to increase after HCMV infection, we monitored ZAP-L and ZAP-S protein levels in WT and control cells at earlier time points (2–10 hpi). Expression levels of ZAP-S were detectable around 6 to 8 hours post HCMV infection and steadily increased over time (Figure 1F).

Taken together, these results show that HCMV infection leads to a strong and steady increase of ZAP-S levels, while ZAP-L is already expressed in uninfected cells but also further induced upon infection and overall stable over a complete cycle of HCMV replication.

### Both ZAP-S and ZAP-L negatively affect HCMV genome replication

To investigate the impact of ZAP on HCMV replication, WT, control, or ZAP KO HFF-1 cells were infected with HCMV and viral genome copies were quantified by qPCR at 1, 3, and 5 days post infection (dpi) (Figure 2A). We included day 1 of infection in our analysis to verify that the knockout of ZAP is not affecting viral entry and that the different cell lines were infected to a similar extent. Indeed, at this early time point when HCMV has not entered the first round of viral genome replication, no significant differences in the number of HCMV genome copies were detected (Figure 2B). At day 3 post infection, when HCMV has completed its first replication cycle, WT and control cells showed significantly lower numbers of viral genome copies compared to ZAP KO cells. At 5 days post infection, HCMV genome copy numbers in WT and control cells were still 5 times lower than in ZAP KO cells (Figure 2B). These results suggest that HCMV genome replication is negatively affected by ZAP-S, ZAP-L, or both.

**Figure 2.**
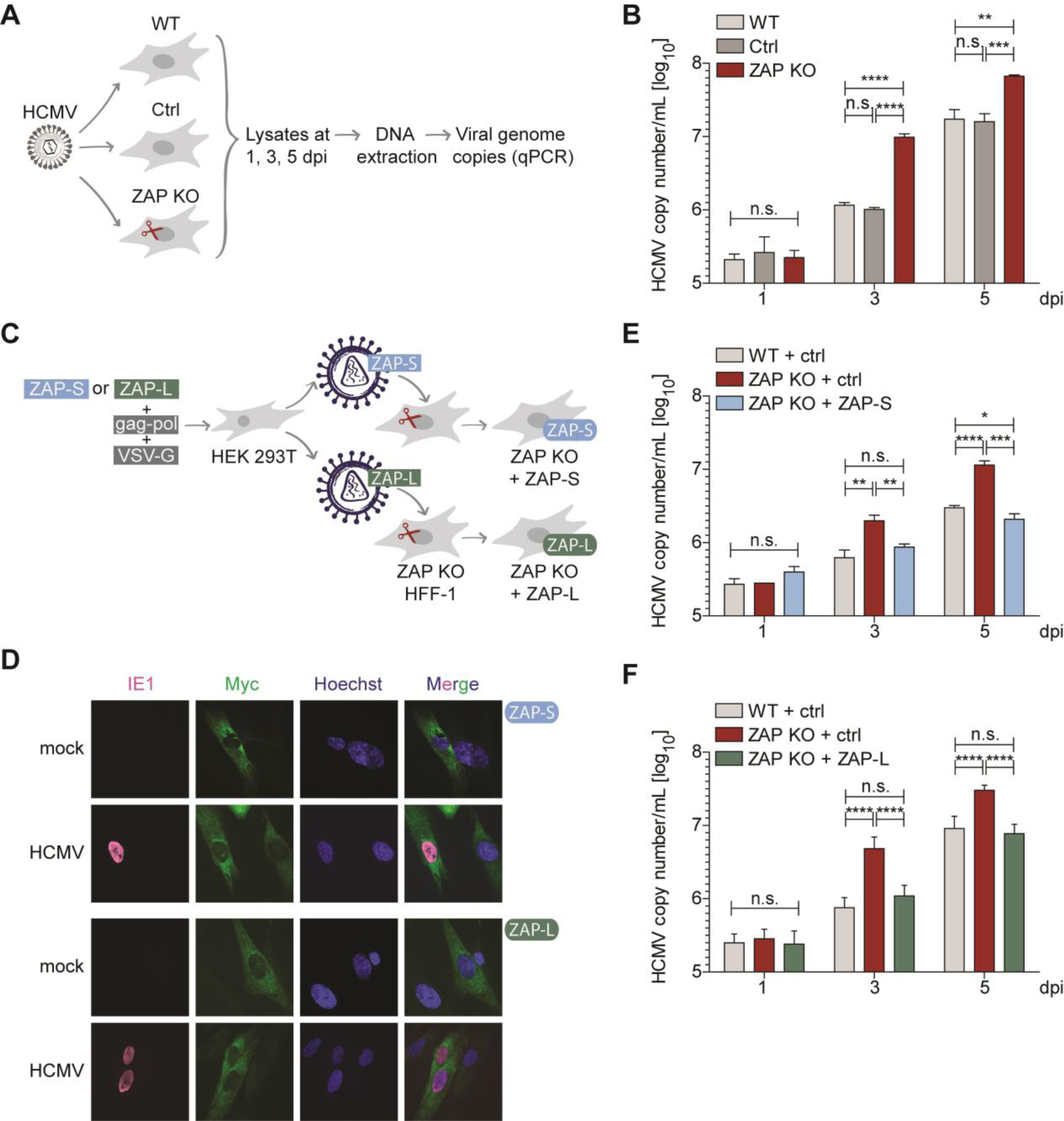
ZAP-S and ZAP-L restrict HCMV replication in HFF-1 cells. **(A)** Schematic representation of the workflow to determine HCMV genome copy numbers. WT, control, or ZAP KO HFF-1 cells were infected with HCMV (MOI 0.1) for 2 hours. Both cells and supernatant were harvested at 1, 3, and 5 days post infection (dpi), followed by DNA extraction and measurement of viral genome copies by qPCR. **(B)** HCMV genome copy numbers from WT, control, or ZAP KO HFF-1 cells were determined as described in **(A)**. HCMV copy numbers/ml are displayed as bar plots showing mean ± S.D. of triplicates. Results shown are one representative of at least three independent experiments using two different ZAP KO cell lines with similar results obtained in all replicates. **(C)** Schematic representation of the workflow to reconstitute ZAP KO HFF-1 cells. HEK 293T cells were transfected with either myc-tagged ZAP-S or ZAP-L expression plasmids together with the packaging (gag-pol) and the envelope (VSV-G) plasmids to produce lentiviruses harbouring ZAP-S or ZAP-L, respectively, followed by transduction of ZAP KO HFF-1 cells. As control, WT and ZAP KO HFF-1 cells were transduced with lentiviruses harbouring empty vector. **(D)** Subcellular localisation of myc-tagged ZAP-S and ZAP-L in ZAP KO HFF-1 cells. ZAP KO cells were transduced as described in **(C)** with either myc-tagged ZAP-S or ZAP-L. Transduced cells were infected by centrifugal enhancement with HCMV (MOI 0.1) and 24 hours post infection cells were fixed for immunolabelling with myc- and IE1-specific antibodies. **(E,F)** HCMV genome copy numbers from WT, ZAP KO, or ZAP KO HFF-1 cells reconstituted with ZAP-S **(E)** or ZAP-L **(F)** were determined as described in **(A)**. HCMV copy numbers/ml are displayed as bar plots showing mean ± S.D. of one **(E)** or two independent **(F)** experiments performed with experimental triplicates. Two independent experiments for both ZAP-S and ZAP-L were performed. Significant changes were calculated using unpaired two-sided Student’s t-tests, n.s. not significant, *\*p* < 0.05, *\*\*p* < 0.01, *\*\*\*p* < 0.001 and *\*\*\*\*p* < 0.0001.

Next, we sought to elucidate which of the two major isoforms contributes to the restriction of HCMV replication. To address this, we reconstituted ZAP KO cells by lentiviral transduction with either an empty vector control, or C-terminally myc-tagged forms of ZAP-S or ZAP-L (Figure 2C). Both ZAP isoforms were codon optimised to avoid recognition and cleavage by the stably expressed gRNA and Cas9 within the KO cell lines. Protein levels of both codon-optimised and WT isoforms were comparable when expressed in HEK 293T cells. Thus, codon optimisation of the ZAP-S or ZAP-L gene did not negatively affect protein expression (Figure EV1). Upon stable expression in ZAP KO HFF-1 cells, both ZAP-S and ZAP-L localised to the cytoplasm under both uninfected and infected conditions (Figure 2D). Next, the reconstituted ZAP KO cells were infected with HCMV and genome copy numbers were analysed as described above (Figure 2A). Strikingly, reconstitution with either ZAP-S (Figure 2E) or ZAP-L (Figure 2F) in ZAP KO cells rescued the phenotype. While HCMV genome copy numbers were equal in infected WT and ZAP-S or ZAP-L reconstituted cells, ZAP KO cells showed significantly higher viral copy numbers. These results show that both ZAP-S and ZAP-L negatively affect HCMV genome replication.

### ZAP negatively affects global expression of early and late HCMV proteins

Given the negative impact of ZAP on HCMV genome replication, we next examined the time course of HCMV infection in the presence or absence of ZAP. Expression of HCMV genes follows a temporal cascade, which begins with the transcription of immediate-early (IE) genes, with the translation of IE proteins starting approximately 6 hours post infection (hpi). IE proteins subsequently transactivate the transcription of early (E) genes, which are mainly involved in viral DNA replication. Early proteins are produced approximately 18-20 hpi and together with a third classical cluster of the so called early-late (E-L) proteins at 48 hpi will mediate the transcription of late (L) genes which code mainly for viral capsid, envelope and tegument components at 72 to 96 hpi (Stinski, 1978; Wathen & Stinski, 1982; Weekes *et al*., 2014). When we monitored the expression levels of the early viral protein UL44 and the late viral protein UL83 in HCMV infected WT and the three ZAP KO cell lines, we observed elevated protein levels of UL44 and UL83 in the absence of ZAP. This suggests that the presence of ZAP negatively affects viral protein expression (Figure 3A), which is in line with our analysis of HCMV genome replication (Figure 2B).

**Figure 3.**
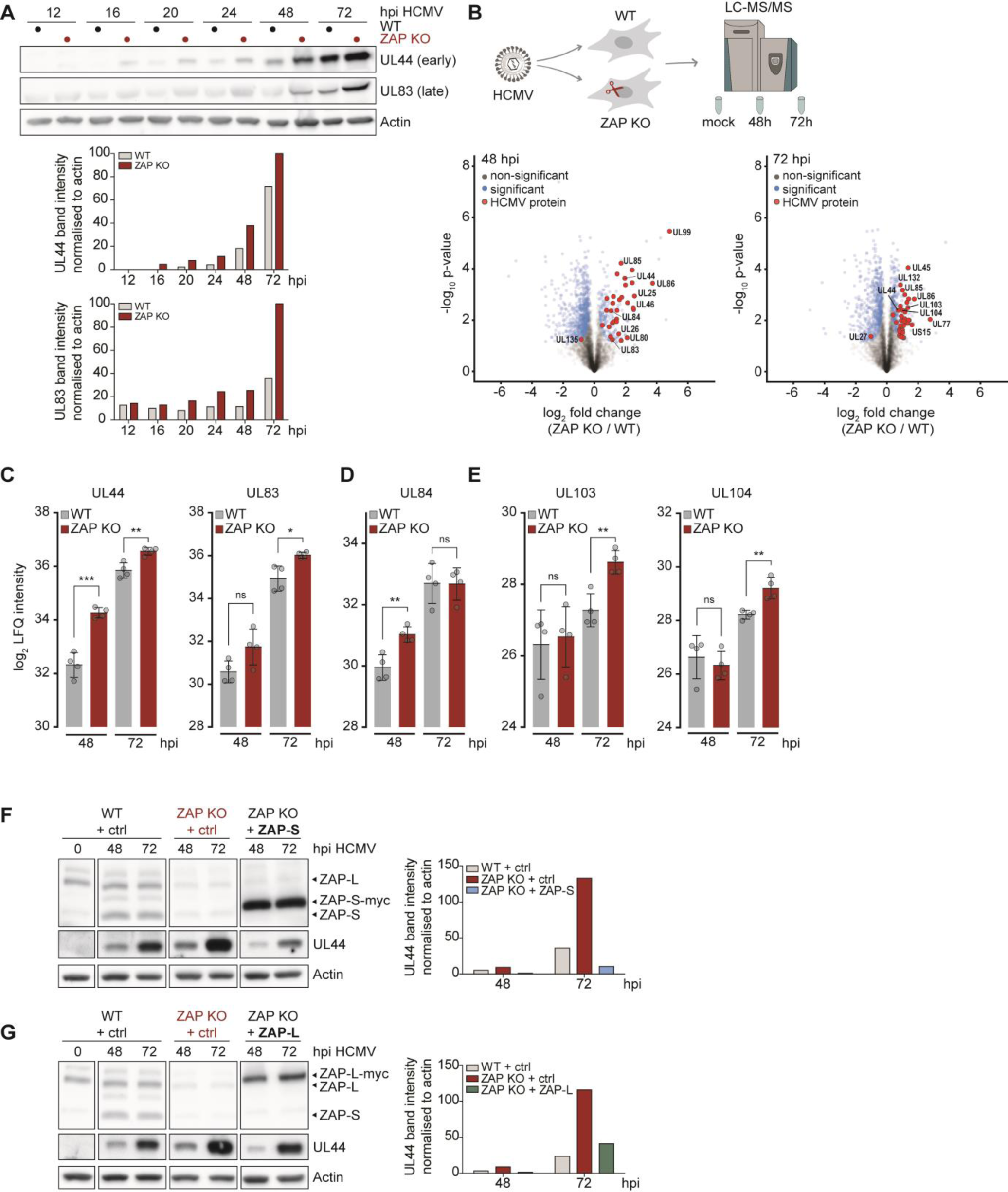
ZAP have a negative impact on early and late HCMV protein levels. **(A)** WT or ZAP KO HFF-1 cells were infected by centrifugal enhancement with HCMV (MOI 0.1) and lysates were analysed at the indicated time points post infection by immunoblotting with specific antibodies against HCMV UL44, HCMV UL83, and actin. One representative experiment performed with three independent ZAP KO cell lines is shown, with similar results in all three experiments. Quantifications of UL44 and UL83 band intensities normalised to actin are represented as bar plots. **(B)** WT and ZAP KO HFF-1 cells were mock treated or infected by centrifugal enhancement with HCMV (MOI 0.1) and cell lysates were subjected to total proteome LC-MS/MS analysis at the indicated time points. Represented are volcano plots (x-axis: log2 fold change, y-axis: -log10 p-value) showing differentially expressed proteins at 48 and 72 hours post HCMV infection with significantly changed proteins (unpaired two-sided Student’s t-test with permutation-based FDR: 0.05, S0=0.1). **(C-E)** Time-resolved expression changes of HCMV UL44 and UL83 **(C)**, UL84 **(D)**, UL103, or UL104 **(E)** in HCMV-infected WT and ZAP KO HFF-1 cells displayed as bar plots showing mean ± S.D. of quadruplicates. **(F, G)** WT, ZAP KO, or ZAP KO HFF-1 cells reconstituted with either ZAP-S **(F)** or ZAP-L **(G)** were infected by centrifugal enhancement with HCMV (MOI 0.1) and lysates were analysed at the indicated time points post infection by immunoblotting with specific antibodies against ZAP, HCMV UL44 and actin. **(F,G)** Quantification of UL44 band intensities normalised to actin is represented as bar plots. One representative of at least 2 independent experiments is shown. Significant changes were calculated using unpaired two-sided Student’s t-tests, n.s. not significant, *\*p* < 0.05, *\*\*p* < 0.01, and *\*\*\*p* < 0.001.

To obtain a global overview of the progression of HCMV infection, we performed whole proteome analyses of WT and ZAP KO HFF-1 cells using liquid chromatography with tandem mass spectrometry (LC-MS/MS) (Table 1). We mock treated or infected WT and ZAP KO HFF-1 cells with HCMV for 48 and 72 hours, thus covering the early-late proteome landscape of viral gene expression. Overall, we observed significantly higher viral protein levels in ZAP KO cells compared to WT cells (Figure 3B). In line with our previous observations (Figure 3A), we detected significantly higher levels of UL44 and UL83 protein in ZAP KO cells (Figure 3C). UL44, considered an early protein, is already expressed at 48 hours, while UL83 protein levels are only increasing at 72 hpi consistent with the kinetics of L gene expression. In line with these observations, the proteome analysis showed other viral proteins that were significantly upregulated in the absence of ZAP. For instance, we detected increased expression of UL84 at 48 hpi, an early protein involved in viral DNA replication (Figure 3D), as well as elevated levels of the late proteins UL103 and UL104 at 72 hpi, similar to UL83 and corresponding to late kinetics (Figure 3E).

Next, we performed reconstitution assays with ZAP-S and ZAP-L as described above (Figure 2C) and analysed UL44 protein expression by immunoblotting (Figure 3F-G). Similar to our analysis of HCMV genome replication, UL44 protein levels in ZAP KO cells reconstituted with either ZAP-S (Figure 3F) or ZAP-L (Figure 3G) were lower than in the absence of ZAP, and comparable to those in WT cells.

Taken together, these results demonstrate that the presence of both main ZAP isoforms negatively affects HCMV protein levels at early and late stages of infection.

### ZAP-S and ZAP-L have a negative impact on early and late HCMV transcripts

Previous studies showed that ZAP directly binds to RNA (Guo *et al*., 2004) and subsequently mediates its degradation by recruiting both the 5’and 3’ RNA degradation machinery (Guo *et al*., 2007; Zhu *et al*., 2011). To decipher whether ZAP affects HCMV mRNA expression, we analysed mRNA levels of the early *UL44* and the late *UL83* transcripts at different stages of the HCMV infection cycle by qRT-PCR in WT and ZAP KO cells. Indeed, in the presence of ZAP, *UL44* and *UL83* mRNA levels were lower (Figure 4A). These results mirror our protein analyses and suggest that ZAP may negatively influence these transcripts by either affecting their expression, stability, or by other, indirect effects. Congruent with our analysis of HCMV protein expression (Figure 3), reconstitution of ZAP KO cells with either ZAP-S or ZAP-L resulted in the rescue of this phenotype (Figure 4B).

**Figure 4.**
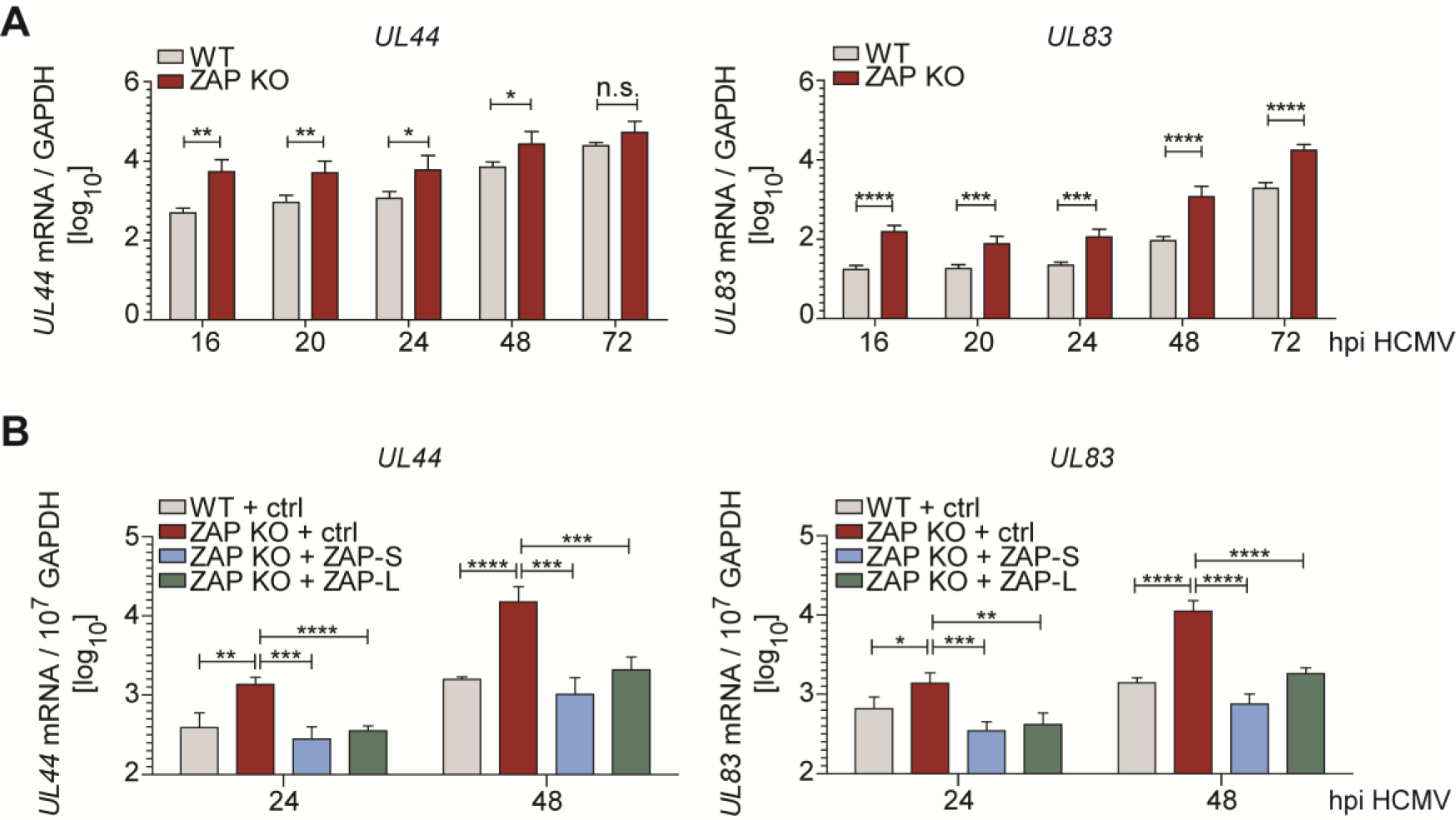
ZAP-S and ZAP-L negatively affect early and late HCMV transcripts. **(A)** WT or ZAP KO HFF-1 cells were infected by centrifugal enhancement with HCMV (MOI 0.1). Total RNA was extracted at indicated time points post infection and mRNA levels of HCMV *UL44* and *UL83* were measured by qRT-PCR. Viral mRNA relative expression (log_10_) normalised to *GAPDH* is displayed as bar plots showing mean ± S.D. of three independent experiments performed with experimental duplicates. Experiments were performed in three independent ZAP KO cell lines and results were combined. **(B)** ZAP KO HFF-1 stably expressing either ZAP-S, ZAP-L, or transduced with empty vector control, and WT cells expressing empty vector, were infected by centrifugal enhancement with HCMV (MOI 0.1). Total RNA was extracted at indicated time points post infection and mRNA levels of HCMV *UL44* and *UL83* were determined by qRT-PCR. Viral mRNA relative expression (log_10_) normalised to *GAPDH* is displayed as bar plots showing mean ± S.D. of two independent experiments performed with experimental duplicates. Significant changes were calculated using unpaired two-sided Student’s t-tests, n.s. not significant, *\*p* < 0.05, *\*\*p* < 0.01, *\*\*\*p* < 0.001 and *\*\*\*\*p* < 0.0001.

Taken together, these results suggest that both ZAP isoforms, ZAP-S and ZAP-L, negatively regulate HCMV mRNA expression.

### ZAP affects stability of early, but not late, HCMV transcripts

Since ZAP was previously described to be involved in mRNA degradation, we studied cellular and viral mRNA stability during HCMV infection of WT and ZAP KO cells. For this, we labelled newly synthesised RNA from 17h-18h and from 71h-72h post infection using 4-thiouridine (4sU) and performed SLAM-seq (thiol-linked alkylation for the metabolic sequencing of RNA) (Herzog *et al*, 2017). Then, we identified the newly synthesised and total RNA using the computational approach GRAND-SLAM (termed Globally refined analysis of newly transcribed RNA and decay rates using SLAM-seq) (Jurges *et al*, 2018) (Figure 5A, Table 2). In agreement with previous studies, we confirmed that *TNFRSF10D* total mRNA was significantly upregulated in uninfected ZAP KO cells, but also expressed higher in the context of HCMV infection (Figure 5B). *TNFRSF10D* encodes for the pro-survival protein TRAIL receptor 4 (TRAILR4, a human cell surface receptor of the TNF-receptor superfamily) and was previously described to be targeted by ZAP at the mRNA level (Todorova *et al*, 2014). Notably, we found another, previously undescribed, anti-apoptotic factor which was significantly upregulated in ZAP KO cells compared to WT cells, *ZMAT3* (encoding for the zinc finger matrin-type protein 3, also known as zinc finger protein Wig-1), with a more pronounced upregulation in the context of HCMV infection (Figure 5B, EV2). For both *TNFRSF10D* and *ZMAT3*, reconstitution with either ZAP-S or ZAP-L rescued these phenotypes as shown by qRT-PCR analyses (Figure EV2). The SLAM-seq results revealed that the upregulation in ZAP KO cells of these two cellular mRNAs in total RNA was not paralleled by the transcription of newly synthesised RNA, where levels were equal between WT and ZAP KO cells (Figure 5B). This shows that in the absence of ZAP, these transcripts have a longer half-life, and indicates that ZAP has an impact on their degradation, but not on their transcription.

**Figure 5.**
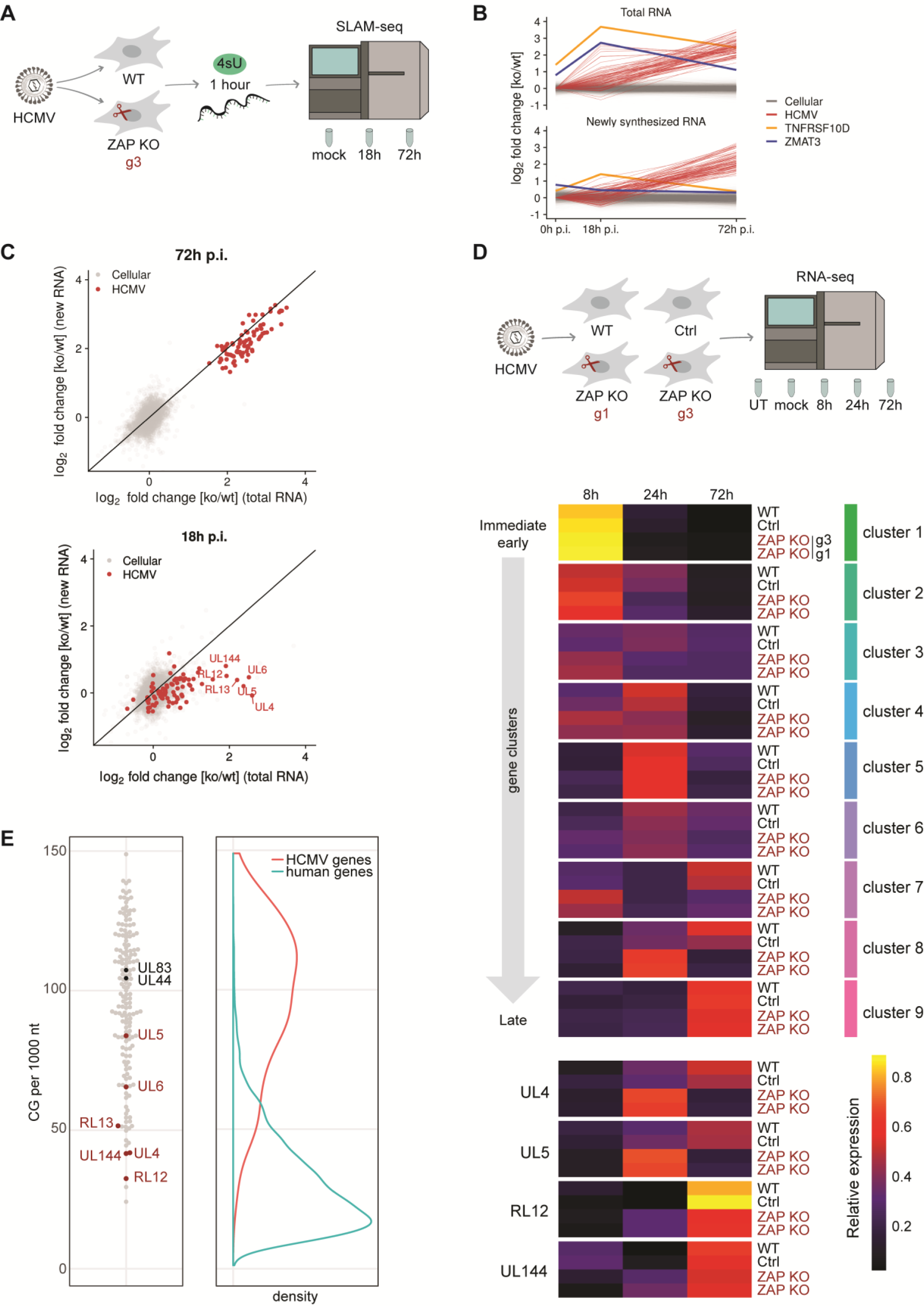
ZAP negatively affects stability of a subset of HCMV transcripts with low CG content. **(A)** WT and ZAP KO HFF-1 cells were untreated or infected by centrifugal enhancement with HCMV (MOI 0.1). Newly synthesised RNA was labelled with 4-thiouridine (4sU) for one hour prior to cell lysis and lysates were taken at 18 and 72 hpi, followed by RNA purification. SLAM-seq was performed to identify newly synthesised and total RNA using GRAND-SLAM. **(B)** Time courses of Log_2_ fold changes of cellular and viral genes (n=6,488) for total (upper panel) and newly synthesised (lower panel) RNA in ZAP KO / WT HFF-1 cells. The values represent the mean of two biological replicates. *TNFRSF10D* and *ZMAT3* are indicated in yellow and purple, respectively. **(C)** Represented are Log_2_ fold changes of cellular and viral genes (n=6,488) for total (x-axis) and newly synthesised RNA (y-axis). Values represent the mean of two biological replicates. **(D)** WT, control, and two independent ZAP KO HFF-1 cell lines were untreated (UT), mock treated, or infected by centrifugal enhancement with HCMV (MOI 0.1). Total RNA was extracted at 8, 24, and 72 hpi, and lysates were subjected to total transcriptome analysis. Relative temporal expression levels of gene clusters and selected individual genes are represented as a heatmap. Expression of HCMV genes was quantified from the RNA-sequencing and relative temporal expression levels calculated by dividing per-sample normalised expression values (fpkm) to the sum of these values from the same gene over all samples/time points. Based on these values, genes were clustered in nine groups representing kinetic classes. Shown are the averages of replicates and clusters (above), as well as averages of replicates of selected individual genes (below). **(E)** For HCMV genes, the CG dinucleotide content per length and gene was calculated from the HCMV TB40/E annotation in accession number MF871618. For human genes, values were calculated from the hg19 Refseq annotation. If multiple transcript isoforms were present in the annotation, values per transcripts were averaged. Values for HCMV genes are shown as a beeswarm plot to the left, and for both human and HCMV as density plots to the right. Selected HCMV genes are labelled in the beeswarm plot.

Regarding viral mRNA transcripts, we observed a 4–12-fold up-regulation of viral genes in ZAP KO cells on both total RNA as well as newly synthesised RNA levels at 72 hpi (Figure 5C). These results indicate that the upregulation on total RNA levels in ZAP KO cells at late times of HCMV infection is predominantly due to increased transcription rates, and not due to an increase in mRNA stability.

Strikingly, at 18 hpi, we identified a subset of viral genes with significantly increased total transcript levels in ZAP KO cells, *UL4*, *UL5*, *UL6*, *RL12*, *RL13*, and *UL144*, which did not correlate with increased *de novo* RNA transcription (Figure 5C), indicating their longer mRNA stability in the absence of ZAP.

In summary, while ZAP does not affect the stability of the majority of viral transcripts, it specifically affects stability of a distinct set of HCMV mRNA transcripts early in infection.

### HCMV infection progresses faster in the absence of ZAP

In order to understand the effect of ZAP on the viral gene expression cascade over time, we analysed the whole transcriptome landscape during HCMV infection by RNA-sequencing (RNA-seq) in WT, control cells, and two independent ZAP KO cell lines (g1 and g3 ZAP KO) (Figure 5D, Table 3). First, we observed that HCMV infection leads to a robust and overall similar induction of ISGs in all cell lines analysed, indicating that the absence of ZAP does not affect the PRR-mediated host response during HCMV infection (Figure EV3). For analysis of the temporal progression of HCMV gene expression, we focused on the mRNA abundance per time point post infection relative to the abundance over all time points. Based on these relative temporal gene expression values in WT cells, genes were clustered into nine groups grouped from immediate-early to late expression, broadly corresponding to a previously published classification (Weekes *et al*., 2014) (Figure 5D, EV4). To simplify the temporal distribution of the viral genes, average relative expression levels of the genes within the clusters were calculated and depicted as a heat map (Figure 5D). Strikingly, the temporal expression patterns of most clusters were shifted to earlier time points in both ZAP KO cell lines, compared to WT and control cells. We observed different subsets of viral genes whose expression was detected earlier in ZAP KO cells compared to WT cells. For instance, ZAP KO cells showed high expression levels of *UL138* or long non-coding *RNA 2.7* (Figure EV4). Notably, these viral transcripts have been previously identified in several studies in relation to the latent cycle of HCMV (Goodrum *et al*, 2007; Rossetto *et al*, 2013; Umashankar *et al*, 2011), a concept that is currently changing and can increasingly be associated with a late-lytic replication programme (Shnayder *et al*, 2018). Differential regulation of the HCMV transcripts *UL4*, *UL5*, *UL6*, *RL12*, *RL13*, and *UL144* identified by SLAM-seq (Figure 5C) was confirmed by RNA-seq, from which selected exemplified transcripts are shown (Figure 5D, lower panel).

In conclusion, ZAP KO cells show an accelerated course of HCMV infection and seem to more rapidly achieve an intracellular environment associated with a late-lytic gene expression programme. These results, in line with our previous findings for viral protein levels, indicate that ZAP delays the progression of the HCMV replication cycle, as reflected in the decelerated course of the viral gene expression cascade.

### ZAP negatively affects a subset of early HCMV transcripts that exhibit CG suppression

Previous studies indicated that ZAP favours CG-dinucleotide enriched RNA sequences and mediates their degradation (Meagher *et al*., 2019; Takata *et al*., 2017). Thus, we compared the CG levels of different viral transcripts to determine whether the HCMV transcripts affected by ZAP are enriched in CG content (Figure 5E). Remarkably, most of the viral transcripts that were significantly upregulated in ZAP KO cells and also showed elevated mRNA stability (Figure 5C-D) exhibit a low CG content that is comparable to that of human genes: *UL4, UL5, UL6, RL12, RL13, and UL144* (Figure 5E). On the contrary, the early *UL44* and the late *UL83* transcripts, whose stability was not affected by ZAP (Figure 5C), have a high CG content (Figure 5E). These results suggest that ZAP may recognise motifs in HCMV transcripts other than CG-dinucleotides.

Taken together, our findings indicate that ZAP may restrict HCMV replication by downregulating the expression of a subset of viral genes with low CG content early during HCMV infection, thus delaying progression of the HCMV infection cycle.

## Discussion

Viral infection induces a virus species specific expression pattern of interferon-stimulated genes (Schoggins *et al*, 2011). To date, more than 300 ISGs have been described, but so far, the function of the majority of the proteins they encode for is poorly understood. ISG function is highly contextual as their effect on viral infection is dependent on the viral entry route, replication mechanism, site of replication and viral assembly, and cell type. Hence, while some ISGs may exert an antiviral activity against some viruses, they may either have a neutral or positive effect on other viruses, or in some instances, be susceptible to viral evasion mechanisms (Gonzalez-Perez *et al*., 2020; Schoggins *et al*., 2014).

HCMV infection leads to the upregulation of a distinct set of cellular proteins during the first 24 hours of infection; 32 of these were classified as ISGs, among them the RNA binding ZAP protein (Weekes *et al*., 2014). While that report did not distinguish between the two major ZAP isoforms ZAP-S and ZAP-L, in this study we delineated their endogenous expression kinetics during HCMV infection. ZAP-L is readily detectable in uninfected cells and its expression slightly increases throughout the first 48 hours of HCMV infection, while ZAP-S protein levels are low in uninfected cells and strongly upregulated from 6 hours post infection onwards. At a late stage of the HCMV life cycle, expression of both ZAP isoforms decreases, which likely reflects the fading type I IFN response rather than HCMV-mediated degradation (Nobre *et al*, 2019; Weekes *et al*., 2014).

Reconstitution of ZAP KO cells with either ZAP-S or ZAP-L showed that both have the potential to restrict HCMV replication to similar levels as endogenous ZAP in wild-type cells. In the course of completing our manuscript, a study was published that reported a negative effect of ZAP on HCMV (Lin *et al*., 2020). The authors analysed expression of four HCMV proteins of three temporal classes, the immediate early proteins 1 (IE1) and 2 (IE2), the early protein UL44 and the late protein UL99. Protein levels of IE2, UL44 and UL99 were reduced in the presence of ZAP, but not that of IE1 (Lin *et al*., 2020). Since ZAP has previously been associated with the binding of CG-rich RNA sequences (Chiu *et al*., 2018; Odon *et al*., 2019; Takata *et al*., 2017), the authors concluded based on a bioinformatics analysis of the CG content of HCMV genes that the low CG content of the *IE1* gene could be an HCMV evasion mechanism to avoid ZAP recognition (Lin *et al*., 2020). While the conclusion drawn by Lin *et al*. (2020) is reasonably supported by their data, the impact of ZAP was only elucidated on four HCMV proteins, without analysing their transcript levels or mRNA stability. Our study took a global approach and examined the whole transcriptome and proteome during HCMV infection in WT and ZAP KO cells. We did not observe an effect of ZAP on total *IE1* transcript levels, confirming the protein expression results by Lin *et al*. (2020). Indeed, we could pinpoint that ZAP strongly delays transcription of the majority of viral genes which consequently results in the delay of viral protein expression.

We reveal, by SLAM-seq, that six HCMV transcripts, namely *UL4*, *UL5*, *UL6*, *RL12*, *RL13,* and *UL144*, have longer half-lives in the absence of ZAP at early times of HCMV infection (18-24 hpi), indicating that these transcripts are destabilised in the presence of ZAP. *UL4*, *UL5*, *UL6*, *RL12*, and *RL13* belong to the RL11 family (Chee *et al*, 1990; Davison *et al*, 2003b). The biological function of the RL11 family members is only poorly understood, but several studies suggest that these membrane-associated proteins may be involved in immune evasion (Atalay *et al*, 2002; Corrales-Aguilar *et al*, 2014; Cortese *et al*, 2012; Davison *et al*, 2003a; Lilley *et al*, 2001). They are largely dispensable for virus growth in cultured fibroblasts, which is often the case for viral proteins involved in immune evasion (Ripalti & Mocarski, 1991; Takekoshi *et al*, 1991). Further, RL13 was described to restrict HCMV growth *in vitro* by an unknown mechanism (Murrell *et al*, 2016; Stanton *et al*, 2010).

The UL144 transmembrane protein has likewise been connected to immune evasion. UL144 is a structural mimic of the tumor necrosis factor receptor superfamily member 14 (TNFRSF14, also known as HVEM for herpesvirus entry mediator) (Benedict *et al*, 1999; Bitra *et al*, 2019; Montgomery *et al*, 1996) and is expressed early in lytic infection and during natural latency in CD14+ monocytes (Benedict *et al*., 1999; Poole *et al*, 2013). Moreover, UL144 is highly variable in sequence between clinical HCMV isolates and some studies reported that certain genotypes can be associated with severe disease outcomes (Benedict *et al*., 1999; Galitska *et al*, 2018; Lurain *et al*, 1999; Waters *et al*, 2010). Its multiple functional consequences on T cell and NK cell-mediated antiviral immunity (Cheung *et al*, 2005; Poole *et al*, 2008; Poole *et al*, 2009; Poole *et al*, 2006; Šedý *et al*, 2013) indicate that UL144 likely plays multiple roles in regulating immunity to HCMV infection during lytic and latent phases of the viral life cycle.

We cannot definitively conclude from our study that the reduced expression of one of these HCMV transcripts or the combination of them is directly responsible for the delayed progression of HCMV infection in ZAP-expressing cells, but given that their function is either poorly understood (RL11 family) or multifactorial (UL144), it can be a plausible explanation. Another possibility is that we are observing two different phenomena in this study: that ZAP on the one hand negatively affects mRNA stability of this specific set of immune evasins, and on the other hand delays viral transcription, presumably in an indirect manner, resulting in reduced viral fitness (Teng *et al*, 2012). Since the function of these immune evasins will be more apparent *in vivo* than *in vitro*, the overall impact of ZAP on HCMV infection can only be fully understood in the clinical setting. This unknown can be addressed, for example, through an analysis of whether specific SNP patterns in ZAP can be predictive of the outcome of HCMV disease.

In addition to viral mRNA transcripts, ZAP has also been described to mediate degradation of specific cellular transcripts (Schwerk *et al*., 2019; Todorova *et al*., 2014). Indeed, we identified two human transcripts whose stability was influenced by ZAP, *TNFRSF10D,* encoding the TRAIL receptor 4 (TRAILR4), and *ZMAT3,* encoding the zinc finger matrin-type protein 3 (also known as zinc finger protein Wig-1). A negative effect of ZAP on *TNFRSF10D* mRNA stability was previously reported, resulting in increased cell sensitivity to TRAIL-mediated apoptosis (Todorova *et al*., 2014). Interestingly, Wig-1 is also a pro-survival factor (Bersani *et al*, 2014).

Altogether, our results demonstrate that ZAP specifically targets host cell transcripts involved in cell survival, as well as a distinct set of HCMV transcripts involved in immune evasion. This raises the question of how ZAP influences mRNA stability of these transcripts - does it bind them directly and mediate their degradation, or does ZAP influence them in an indirect manner? Several studies describe the importance of CG-rich sequences for binding of ZAP to RNA, which eventually leads to the degradation of this RNA (Chiu *et al*., 2018; Luo *et al*, 2020; Meagher *et al*., 2019; Odon *et al*., 2019; Takata *et al*., 2017). In comparison to the human genome, which has low CG content (Antequera & Bird, 1993; Bestor & Coxon, 1993), the HCMV genome presents the highest CG content among human *Betaherpesvirinae* (Sharma *et al*, 2016), which makes HCMV transcripts a good target for ZAP-mediated degradation. However, in our study, the HCMV transcripts whose stability was affected by ZAP display low CG content, while the stability of HCMV transcripts with high CG content such as *UL44* or *UL83* and the majority of other viral transcripts were not affected. In addition, ZAP also targets host mRNAs with low CG content. Hence, we propose that ZAP may bind transcripts via motifs other than CG-rich sequences, e.g. AU-rich elements, which can be bound by ZAP (Odon *et al*., 2019; Schwerk *et al*., 2019), or via more specific motifs that have yet to be identified. CLIP-seq assays are challenging to perform in the context of infection, but are a desirable approach to prove direct interaction between ZAP and viral transcripts.

While both ZAP isoforms, ZAP-S and ZAP-L, could restrict HCMV infection in our study, recent studies propose diverse functions for one or the other isoform during infection with different viruses. This is the case for Sindbis virus (SINV), an RNA alphavirus previously shown to be inhibited by ZAP (Bick *et al*., 2003; Schwerk *et al*., 2019), and the DNA virus Modified Vaccinia Virus Ankara (MVA) (Peng *et al*., 2020). Schwerk and colleagues observed that ZAP-L, which can be farnesylated at its C-terminus (which is lacking in ZAP-S) and thereby be targeted to membranes (Charron *et al*., 2013), colocalises with SINV RNA intermediates in distinct foci in the cytoplasm (Schwerk *et al*., 2019). Similarly, for MVA, which also replicates in the cytoplasm, no impact of ZAP on viral transcription was observed, but rather an effect of ZAP-L on viral assembly (Peng *et al*., 2020). Nonetheless, for HCMV, which replicates its DNA genome in the nucleus, we did not observe distinct foci in ZAP-L expressing cells, but a similar diffuse cytoplasmic localisation for both

ZAP-L and ZAP-S in uninfected and infected HFF-1 cells. Hence, in the case of the DNA virus HCMV, ZAP-L and ZAP-S may exert redundant functions, both targeting mRNAs localised in the cytoplasm. Interestingly, ZAP-S was described to act as a negative feedback regulator of the IFN response later in infection with SINV by destabilising the IFNβ transcript (Schwerk *et al*., 2019). However, we did not observe an effect for ZAP-S on IFN signalling pathways, which likely corresponds with HCMV encoding multiple viral evasion proteins that downmodulate the IFN response (Gonzalez-Perez *et al*., 2020; Stempel *et al*, 2019). Hence, a possible effect of ZAP-S in this regard may be overshadowed in the context of infection with this complex herpesvirus.

Altogether, these findings show the multiple layers of complexity of this RNA binding protein, highlighting its different facets depending on the virus species it encounters. For HCMV, ZAP appears on the scene at early time points of infection, decelerating the viral gene expression cascade, and handpicks for degradation a distinct set of viral transcripts encoding for viral immune evasins, illustrating its potent role as an antiviral restriction factor for this complex herpesvirus.

## Materials and methods

### Cell lines

Primary human foreskin fibroblasts (HFF-1; SCRC-1041), MRC-5 (CCL-171) and human embryonic kidney 293T cells (HEK 293T; CRL-3216) were obtained from ATCC. HEK 293T and MRC-5 cells were maintained in Dulbecco’s modified Eagle’s medium (DMEM; high glucose) supplemented with 8% fetal calf serum (FCS), 2 mM glutamine (Gln), and 1% penicillin/streptomycin (P/S). HFF-1 cells were maintained in DMEM (high glucose) supplemented with 15% FCS, 1% P/S and 1% non-essential amino acids (NEAA). Cells were cultured at 37°C and 7.5% CO_2_.

### Viruses

The wild-type HCMV TB40-BAC4 (hereinafter designated as HCMV WT) was characterised previously (Sinzger *et al*, 2008) and kindly provided by Martin Messerle (Institute of Virology, Hannover Medical School, Germany). HCMV BACs were reconstituted after transfection of MRC5 cells with purified BAC DNA. Reconstituted virus was propagated in HFF-1 cells and virus was purified on a 10% Nycodenz cushion. The resulting virus pellets were resuspended in virus standard buffer (50 mM Tris-HCl pH 7.8, 12 mM KCl, 5 mM EDTA) and stored at -70°C. Infectious titre was determined by standard plaque assay and IE1 labelling using HFF-1.

### Plasmids

Expression plasmids for Firefly Luciferase (FFLuc, control) and ZAP-S (short isoform of ZAP) in pTRIP-IRES-RFP as well as pCMV-VSV-G and pCMV-gag/pol plasmids were described previously (Schoggins *et al*., 2011) and kindly provided by John Schoggins (University of Texas Southwestern Medical Center, Dallas, Texas). pcDNA4-HA-ZAP-L (long isoform of ZAP) (Kerns *et al*., 2008) was kindly provided by Chad Swanson (Department of Infectious Diseases, School of Immunology and Microbial Sciences, King’s College London). ZAP-S and ZAP-L were subcloned into pEF1-V5/His (Thermo Fisher Scientific) via the *KpnI*/*XbaI* sites to generate pEF1-ZAP-S-V5/His and pEF1-ZAP-L-V5/His, respectively. Exchange of V5/His to myc/His was performed using the Q5 site-directed mutagenesis kit (NEB #E0554) according to the manufacturer’s protocol resulting in pEF1-ZAP-S-myc/His and pEF1-ZAP-L-myc/His. In order to reconstitute ZAP-S and ZAP-L expression in ZAP KO cell lines, codon optimisation of the ZAP-S and ZAP-L coding region was performed to prevent binding of the constitutively expressed gRNA and Cas9. For this, nucleotides 103-219 of the ZAP-S and ZAP-L coding region (spanning across the binding sites for gRNA 1 and gRNA 3, see Figure 1C) were codon optimised using the Q5 site-directed mutagenesis kit, resulting in pEF1-ZAP-S-myc/His and pEF1-ZAP-L-myc/His codon-optimised. A pTRIP-IRES-RFP empty vector was generated by replacing the coding region of ZAP-S from pTRIP-IRES-RFP ZAP-S (received from John Schoggins) by the multiple cloning site of pWPI vectors to obtain *PmeI*, *SdaI*, *SgsI*, *BamHI*, *XmaI*, *RgaI* and *XhoI* restriction sites for further subcloning. Codon-optimised versions of ZAP-S and ZAP-L were subcloned into the newly generated pTRIP-IRES-RFP empty vector via the *SgsI*/*BamHI* restriction sites to generate pTRIP-IRES-RFP ZAP-S-opt-myc/His and pTRIP-IRES-RFP ZAP-L-opt-myc/His. All constructs were verified by sequencing. Oligo sequences as well as sequences of all constructs are available upon request.

The expression plasmid for gRNA cloning and CRISPR/Cas9-mediated gene editing, pLK05.U6.sgRNA(BsmBI,stuffer).EFS.SpCas9.P2A.tagRFP (Heckl *et al*, 2014), was kindly provided by Dirk Heckl (Experimental Pediatrics, Hannover Medical School, Germany). The corresponding envelope and packaging plasmids pMD2.G and psPAX2 were purchased from AddGene (#12259 and #12260, respectively).

### Antibodies and reagents

Mouse monoclonal anti-pp65 (#ab6503, clone 3A12) was obtained from Abcam and mouse monoclonal anti-ICP36 (anti-UL44) (#MBS530793, clone M612460) was purchased from MyBioSource. Mouse monoclonal anti-hZAP (ZC3HAV1) (#66413-1-Ig, clone 1G10B9) and rabbit polyclonal anti-hZAP (#16820-1-AP) were obtained from Proteintech. Mouse monoclonal anti-actin (A5441, clone AC-15) was obtained from Sigma-Aldrich. Rabbit monoclonal anti-myc (#2278, clone 71D10) was obtained from Cell Signaling. Mouse monoclonal anti-IE1 (clone 63-27, originally described in Andreoni *et al*, 1989) was a kind gift from Jens von Einem (Institute of Virology, Ulm University Medical Center, Ulm). Alexa Fluor^®^-conjugated secondary antibodies were purchased from Invitrogen. The transfection reagent Lipofectamine 2000 was purchased from Life Technologies, Polybrene was obtained from SantaCruz. OptiMEM was purchased from Thermo Fisher Scientific. Protease inhibitors (4693116001) were purchased from Roche. Recombinant human IFNβ was purchased from PeproTech (#300-02BC).

### Generation of ZAP knockout cells using CRISPR/Cas9-mediated genome editing

Custom gRNAs targeting the first exon of the ZAP coding region, thus disrupting expression of ZAP-S and ZAP-L, were designed using CRISPOR software (http://crispor.tefor.net) (Haeussler *et al*, 2016) and cloned into the lentiviral pLKO5 vector (kindly provided by Dirk Heckl, Martin-Luther-University in Halle, Germany). The pLKO5 vector constitutively expresses the introduced gRNA under the control of a U6 promoter. SpCas9 with a P2A cleavage site followed by RFP is under the control of the EF1α short promoter and results in the constitutive expression of SpCas9 and an RFP reporter for cell sorting. Three different gRNAs targeting the ZAP coding region and a non-targeting control gRNA were generated and cloned into the pLKO5 vector via the *BsmBI* restriction site. ZAP gRNA 1 targets exon 1 at nucleotide 149, ZAP gRNA2 at nucleotide 53 and ZAP gRNA3 at nucleotide 191. The gRNA sequences are as follows: ZAP-g1_FOR: 5’-CACCGGCCGGGCCCGACCGCTTTG; ZAP-g1_REV:

5’-AAACCAAAGCGGTCGGGCCCGGCC; ZAP-g2_FOR:

5’-CACCGCAAAATCCTGTGCGCCCACG; ZAP-g2_REV:

5’-AAACCGTGGGCGCACAGGATTTTGC; ZAP-g3_FOR:

5’-CACCGGCCGGGATCACCCGATCGG; ZAP-g3_REV:

5’-AAACCCGATCGGGTGATCCCGGCC; control-gRNA_FOR:

5’-CACCGGATTCTAAAACGGATTACC; control-gRNA_REV:

5’-AAACGGTAATCCGTTTTAGAATCC. For lentivirus production, HEK 293T cells (730,000 cells per well, 6-well format) were transfected with 400 ng pMD2.G, 1,600 ng psPAX2 and 2,000 ng pLKO5 plasmid (containing the respective gRNA) complexed with Lipofectamine. 16 hours post transfection, medium was changed to lentivirus harvest medium (DMEM h.gl. supplemented with 20% FCS, 1% P/S and 10mM HEPES). 48 hours post transfection, lentivirus was harvested, diluted 1:2 with HFF-1 medium, and polybrene was added to a final concentration of 4 µg/ml. HFF-1 cells were seeded the day before transduction in a 6-well format with 250,000 cells/well. For transduction, HFF-1 medium was replaced by medium containing lentivirus and cells were transduced by centrifugal enhancement at 684 x g and 30°C for 90 minutes. 3 hours post transduction, medium was replenished to fresh HFF-1 medium. Successfully transduced cells were sorted by flow cytometry for RFP signal 72 hours post transduction to obtain a cell population devoid of ZAP expression. Cas9-mediated knockout of ZAP was verified by immunoblot.

### Reconstitution assays

For reconstitution of ZAP-S or ZAP-L expression in ZAP KO cell lines, lentiviral transduction was performed as described above. Briefly, 2,000 ng pTRIP-IRES-RFP empty vector or pTRIP-IRES-RFP containing codon-optimised C-terminally myc-tagged ZAP-S or ZAP-L together with 400 ng pCMV-VSV-G and 1,600 ng pCMV-gag/pol complexed with Lipofectamine were transfected into HEK 293T cells and medium was changed to lentivirus harvest medium the next day. 48 hours post transfection, WT or the indicated ZAP KO HFF-1 cells were lentivirally transduced. 72 hours post transduction, cells were counted and 100,000 cells per well were seeded in a 24-well format. The next day, cells were infected with HCMV WT at an MOI of 0.1 and infection was enhanced by centrifugation at 684 x g for 45 min at 30°C. After centrifugation, cells were incubated at 37°C for 30 min followed by replacement of virus-containing medium with fresh HFF-1 medium. At indicated time points post infection, cells were lysed for analysis by immunoblot or qRT-PCR as described below.

### Immunoblotting

For the analysis of viral protein kinetics upon HCMV infection, HFF-1 WT or ZAP KD cells (100,000 cells/well in a 24-well format) were infected with HCMV WT at an MOI of 0.1 and the infection was enhanced by centrifugation at 684 x g at 30°C for 45 min. The moment when the virus was added to the cells was defined as time point 0. After centrifugation, cells were incubated at 37°C for 30 minutes followed by replacement of virus-containing medium with fresh HFF-1 medium. Cells were lysed at indicated time points using radioimmunoprecipitation (RIPA) buffer (20 mM Tris–HCl pH 7.5, 1 mM EDTA, 100 mM NaCl, 1% Triton X-100, 0.5% sodium deoxycholate, 0.1% SDS). Protease inhibitors were added freshly to all lysis buffers prior to use. Cell lysates and samples were separated by SDS–PAGE and transferred to PVDF membrane (GE Healthcare) using wet transfer and Towbin blotting buffer (25 mM Tris, 192 mM glycine, 20% (v/v) methanol). Membranes were probed with the indicated primary antibodies and respective secondary HRP-coupled antibodies diluted in 5% w/v non-fat dry milk or 5% BSA in TBS-T. Immunoblots were developed using SuperSignal West Pico (Thermo Fisher Scientific) chemiluminescence substrates. Membranes were imaged with a ChemoStar ECL Imager (INTAS) and quantified using the LabImage 1D software (INTAS).

### Immunofluorescence

ZAP KO HFF-1 cells were lentivirally transduced as described above. Briefly, 2,000 ng pTRIP-IRES-RFP containing codon-optimised C-terminally myc-tagged ZAP-S or ZAP-L together with 400 ng pCMV-VSV-G and 1,600 ng pCMV-gag/pol complexed with Lipofectamine were transfected into HEK 293T cells and medium was changed to lentivirus harvest medium the next day. 48 hours post transfection, ZAP KO HFF-1 cells were lentivirally transduced. 72 hours post transduction, cells were counted and 20,000 cells per well were seeded in a µ-Slide 8 Well (ibidi #80826).The next day, cells were mock infected or infected with HCMV WT at an MOI of 0.1 and infection was enhanced by centrifugation at 684 x g for 45 min at 30°C. After centrifugation, cells were incubated at 37°C for 30 min followed by replacement of virus-containing medium with fresh HFF-1 medium. Cells were fixed 24 hpi using 4% PFA in PBS for 20 min at room temperature. Cells were washed three times with PBS, followed by permeabilization using 0.4% Triton X-100 in PBS for 10 min at room temperature. Cells were washed three times with PBS and blocked with 4% BSA in PBS for 45 min. Cells were stained with the indicated primary antibodies and respective secondary antibodies coupled to Alexa488, or Alexa647, and Hoechst (Thermo Fisher Scientific, #33342) diluted in 4% BSA in PBS for 45 min at room temperature in the dark. Imaging was done on a Nikon ECLIPSE Ti-E-inverted microscope equipped with a spinning disk device (Perkin Elmer Ultraview), and images were processed using Volocity software (version 6.2.1, Perkin Elmer).

### Quantitative RT-PCR

HFF-1 WT or ZAP KO cells were infected with HCMV WT as described above. Cells were lysed in RLT buffer and RNA was purified using the Jena Analytik RNA isolation kit (845-KS-2040250), following the manufacturer’s protocol. After RNA extraction, 1,500 ng of RNA per sample was used for further processing. DNase treatment and cDNA synthesis was performed with the iScript gDNA clear kit (172-5035) following the manufacture’s protocol. Generated cDNA was diluted 1:5 before performing qPCR to obtain 100 µl of cDNA. For quantification of gene transcripts, 5 µl of cDNA per sample were used and qRT-PCR was performed using the GoTaq^®^ qPCR Master Mix (Promega, A6001) on a LightCycler 96 (Roche). GAPDH was used for normalisation. The following oligo sequences were used: GAPDH_FOR: 5’-GAAGGTGAAGGTCGGAGTC; GAPDH_REV: 5’-GAAGATGGTGATGGGATTTC; UL44_FOR: 5’-CGCGACGTTACTTTGATTTGAG; UL44_REV:

5’-ATTCGGACGCCGACATTAG; UL83_FOR: 5’-AACCAAGATGCAGGTGATAGG; UL83_REV:

5’-AGCGTGACGTGCATAAAGA; TNFRSF10D_FOR: 5’-CTGCTGGTTCCAGTGAATGACG;

TNFRSF10D_REV: 5’-TTTTCGGAGCCCACCAGTTGGT; ZMAT3_FOR: 5’-GCTCTGTGATGCCTCCTTCAGT; ZMAT3_REV: 5’-TTGACCCAGCTCTGAGGATTCC.

### Determination of HCMV genome copy numbers

HFF-1 WT, control, or ZAP KO cells were infected with HCMV WT as described above. At indicated time points, cells were scraped into the supernatants and cells and supernatant were harvested together. DNA from 200 µl of the samples was extracted using the Qiagen DNeasy^®^ Blood & Tissue kit (#69504) following the manufacturer’s protocol. Extracted DNA was diluted 1:10 prior to qPCR for the analysis of HCMV genome copy numbers. HCMV DNA copy numbers were quantified with a real time quantitative PCR as described previously (Henke-Gendo *et al*, 2012). Copy numbers were harmonized to the 1^st^ WHO International Standard for Human Cytomegalovirus for Nucleic Acid Amplification Techniques (NIBSC # 09/162).

### Total transcriptome analyses (RNA sequencing)

HFF-1 WT, control and two independent ZAP KO cells (250,000 cells/well in a 6 well-plate format) were infected by centrifugal enhancement at 684 x g and 30°C for 45 minutes with HCMV WT at an MOI of 0.1. The moment when the virus was added to the cells was defined as time point 0. After centrifugation, cells were incubated at 37°C for 30 minutes followed by removal of the virus-containing medium and washed with fresh DMEM once. Medium was then replaced with previously conditioned medium (medium cells were originally seeded in, kept at 37°C during the infection time). Cells were lysed at the indicated time points using Trizol for 2 minutes at room temperature and kept at -70°C. Two wells were combined to obtain around 500,000 cells per sample. Total RNA was isolated using the RNA clean and concentrator kit (Zymo Research), according to the manufacturer’s instructions. Sequencing libraries were prepared using the NEBNext Ultra II Directional RNA Library Prep Kit for Illumina (NEB, cat #E7760) following polyA RNA enrichment (NEB cat #E7490) with 9 cycles PCR amplification, and sequenced on a HiSeq 4000 1x50 cycles flowcell.

Alignments were done using hisat2 (Kim *et al*, 2015). Sequencing reads were aligned to the hg19 version of the human genome using standard parameter, using the Refseq gtf file downloaded from the UCSC genome browser. Reads were then quantified using quasR (Gaidatzis *et al*, 2015) and the above mentioned gtf file, or the HCMV TB40/E annotation (accession number MF871618). Differential expression and corresponding p-values were calculated using edgeR (McCarthy *et al*, 2012). Plots were created using ggplot2 (Wickham, 2009) and pheatmap v.1.0.12 (Kolde, R. 2019. pheatmap: Pretty Heatmaps. https://cran.r-project.org/web/packages/pheatmap/index.html).

### SLAM sequencing

HFF-1 WT or ZAP KO (g3) cells (250,000 cells/well in a 6-well format) were untreated or infected with HCMV WT at an MOI of 0.1 and the infection was enhanced by centrifugation at 684 x g at 30°C for 45 min. The moment when the virus was added to the cells was defined as time point 0. After centrifugation, cells were incubated at 37°C for 30 minutes followed by removal of the virus-containing medium, one wash with DMEM and replacement with fresh medium. Newly synthesised RNA was labelled with 4-thiouridine (4sU) for one hour prior to cell lysis. Cells were lysed at the indicated time points using Trizol for 5 minutes at room temperature and kept at -70°C. Two wells were combined to obtain around 500,000 cells per sample. Total RNA was isolated using the DirectZOL kit (Zymo Research) according to the manufacturer’s instruction including the optional on-column DNAse digestion. The 4sU alkylation reaction was essentially performed as published before (Herzog *et al*., 2017). Briefly, 7.5 – 15 µg total RNA were incubated in 1x PBS (pH 8) containing 50% DMSO and 10 mM IAA at 50°C for 15 minutes. The reaction was quenched with 100 mM DTT and RNA purified using the RNeasy kit (Qiagen). Quality and integrity of total RNA was controlled on 5200 Fragment Analyzer System. The RNA sequencing library was generated from 100 ng total RNA using NEBNext® Single Cell/Low Input RNA Library to manufacturés protocols. The libraries were sequenced on Illumina NovaSeq 6000 using NovaSeq 6000 S1 Reagent Kit (300 cycles, paired end run 2x 150 bp) with an average of 40 x 10^6^ reads per RNA sample.

SLAM-seq was performed in duplicates to identify newly synthesised and total RNA using GRAND-SLAM. Sequencing adapters (AGATCGGAAGAGCACACGTCTGAACTCCAGTCA, AGATCGGAAGAGCGTCGTGTAGGGAAAGAGTGT) were trimmed using Trimmomatic (v0.39) (Bolger *et al*, 2014). Reads were mapped to a combined index of the human genome (Hg38 / Ensembl v90) and the HCMV genome (accession number KF297339.1) using STAR (v2.5.3a) with parameters --outFilterMismatchNmax 20 --outFilterScoreMinOverLread 0.3 --outFilterMatchNminOverLread 0.3 --alignEndsType Extend5pOfReads12 --outSAMattributes nM MD NH. We used GRAND-SLAM (v2.0.5d) (Jurges *et al*., 2018) to estimate the new-to-total RNA ratios. We only used the parts of the reads that were sequenced by both mates in a read pair (parameter -double) for the estimation. New RNA was computed by multiplying total RNA with the maximum *a posteriori* estimate of the new-to-total RNA ratio. For further analyses, we removed all cellular genes that had less than 10 transcripts per million transcripts (TPM) in more than 6 (cellular genes) or 2 (viral genes) samples. To remove artefacts due to imprecise quantification, we furthermore removed all viral genes with less than 100 new reads. Log2 fold changes were estimated using PsiLFC (Erhard, 2018) with uninformative prior (corresponding to no pseudocounts). Normalization factors were computed from total RNA such that the median log2 fold change was 0 and applied to both total and new RNA.

### Total proteome analyses using LC-MS/MS

HFF-1 WT or ZAP KO (250,000 cells/well in a 6-well format) were infected with HCMV WT at an MOI of 0.1 and the infection was enhanced by centrifugation at 684 x g at 30°C for 45 min. The moment when the virus was added to the cells was defined as time point 0. After centrifugation, cells were incubated at 37°C for 30 minutes followed by removal of the virus-containing medium, one wash with DMEM and replacement with fresh medium. At indicated time points, cells were washed with PBS once and then collected in 300 µl of fresh PBS per well. Cell pellets were frozen at -70°C. Two wells were combined to obtain a total of 500,000 cells in total per condition. Quadruplicates of HCMV infected HFF-1 cells were analysed at 48 and 72 hours post infection. For each replicate, cells were washed with PBS, lysed in SDS lysis buffer (4% SDS, 10 mM DTT, 50 mM Tris/HCl pH 7.6), boiled at 95°C for 5 min and sonicated (4°C, 10 min, 30 sec on, 30 sec off; Bioruptor). Protein concentrations of cleared lysates were normalised and cysteines were alkylated with 5.5 mM IAA (20 min, 25°C, in the dark). SDS was removed by protein precipitation with 80% (v/v) acetone (-20°C, overnight), protein pellets were washed with 80% (v/v) acetone and resuspended in 40 µl U/T buffer (6 M urea, 2 M thiourea in 10 mM HEPES, pH 8.0). Protein digestion was performed by subsequent addition of 1 µg LysC (3h, 25°C) and 1 µg Trypsin in 160 µl digestion buffer (50 mM ammonium bicarbonate, pH 8.0) at 25°C overnight. Peptides were desalted and concentrated using C18 Stage-Tips as described previously (Hubel *et al*, 2019). Purified peptides were loaded onto a 50 cm reverse-phase analytical column (75 µm diameter; ReproSil-Pur C18-AQ 1.9 µm resin; Dr. Maisch) and separated using an EASY-nLC 1200 system (Thermo Fisher Scientific). A binary buffer system consisting of buffer A (0.1% formic acid in H2O) and buffer B (80% acetonitrile, 0.1% formic acid in H2O) with a 120 min gradient [5-30% buffer B (95 min), 30-95% buffer B (10 min), wash out at 95% buffer B (5 min), decreased to 5% buffer B (5 min), and 5% buffer B (5 min)] was used at a flow rate of 300 nl per min. Eluting peptides were directly analysed on a Q-Exactive HF mass spectrometer (Thermo Fisher Scientific). Data-dependent acquisition included repeating cycles of one MS1 full scan (300–1,650 m/z, R = 60,000 at 200 m/z) at an ion target of 3 x 10^6^, followed by 15 MS2 scans of the highest abundant isolated and higher-energy collisional dissociation (HCD) fragmented peptide precursors (R = 15,000 at 200 m/z). For MS2 scans, collection of isolated peptide precursors was limited by an ion target of 1 x 10^5^ and a maximum injection time of 25 ms. Isolation and fragmentation of the same peptide precursor was eliminated by dynamic exclusion for 20 s. The isolation window of the quadrupole was set to 1.4 m/z and HCD was set to a normalised collision energy of 27%. Raw files were processed with MaxQuant (version 1.6.14.0) using the standard settings and label-free quantification (LFQ) and match between runs options enabled. Spectra were searched against forward and reverse sequences of the reviewed human proteome including isoforms (UniprotKB, release 01.2019) and of the HCMV proteins by the built-in Andromeda search engine (Tyanova *et al*, 2016a).

### Statistical analyses

The output of MaxQuant was analysed with Perseus (version 1.6.14.0, Tyanova *et al*, 2016b), R (version 3.6.0), RStudio (version 1.2.1335) and GraphPad Prism (version 7.04). Detected protein groups identified as known contaminants, reverse sequence matches, only identified by site or quantified in less than 3 out of 4 replicates in at least one condition were excluded. Following log2 transformation, missing values were imputed for each replicate individually by sampling values from a normal distribution calculated from the original data distribution (width = 0.3*s.d., downshift = -1.8*s.d.). Differentially expressed protein groups between biological conditions were identified via two-sided Student’s T-tests corrected for multiple hypotheses testing applying a permutation-based FDR (250 randomizations).

For qRT-PCR and genome copy number quantification, differences between data sets were evaluated after log transformation by Student’s *t*-test (unpaired, two-tailed), using GraphPad Prism version 5.0 (GraphPad Software, San Diego, CA). *P* values < 0.05 were considered statistically significant.

## Acknowledgements

We would like to thank Niels Lemmermann and Baca Chan for their insightful comments as well as John Schoggins for generously providing us with ISG expression plasmids. We also thank Georg Wolf and Christine Standfuβ-Gabisch for excellent technical assistance.

The PhD scholarship to ACGP was funded by the European Union’s Horizon 2020 research and innovation programme (H2020) under the Marie Skłodowska-Curie Innovative Training Networks Programme MSCA-ITN GA 675278 EDGE (Training Network providing cutting-EDGE knowlEDGE on Herpes Virology and Immunology). MS was funded by the Deutsche Forschungsgemeinschaft (DFG, German Research Foundation) – Project number 158989968 - SFB 900, and MMB by the SMART BIOTECS alliance between the Technische Universität Braunschweig and the Leibniz Universität Hannover, an initiative supported by the Ministry of Science and Culture (MWK) of Lower Saxony, Germany, and the Helmholtz Association (W2/W3-090).

## Author contributions

Conceptualization, ACGP, MS, MMB; Methodology, ACGP, MS, EW, A Piras, TH; Investigation, ACGP, MS, EW, CU, FH, MMB; Writing-Original Draft, ACGP, MS, MMB; Writing-Review & Editing, ACGP, MS, EW, CU, A Piras, TH, AH, ML, A Pichlmair, FE, LD, MMB; Funding Acquisition, MMB; Resources, AH, ML, A Pichlmair, LD, MMB; Supervision, MS, MMB.

## Tables and their legends

Table 1. Proteome analyses.

Table 2. SLAM-sequencing.

Table 3. Transcriptome analyses.

## Expanded view figure legends

**Figure EV1.**
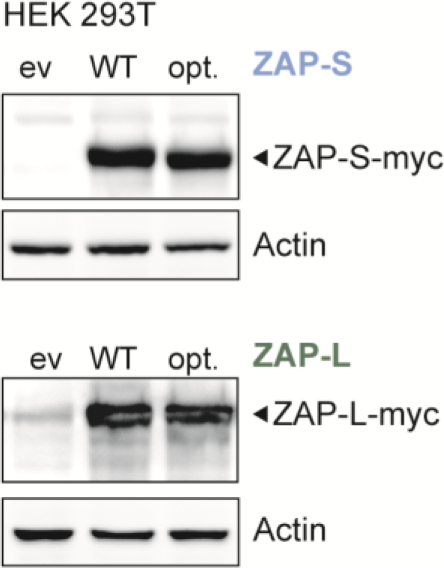
Codon-optimisation of ZAP does not affect ZAP protein levels. HEK 293T cells were transfected with either pEF empty vector (ev), pEF1-ZAP-S-myc/His (WT), or pEF1-ZAP-S-myc/His codon-optimised (opt.) (upper panel) or with pEF1-ZAP-L-myc/His WT or codon-optimised (opt.) expression constructs (lower panel). Expression levels of ZAP were determined by immunoblotting using a ZAP-specific antibody. Actin served as loading control.

**Figure EV2.**
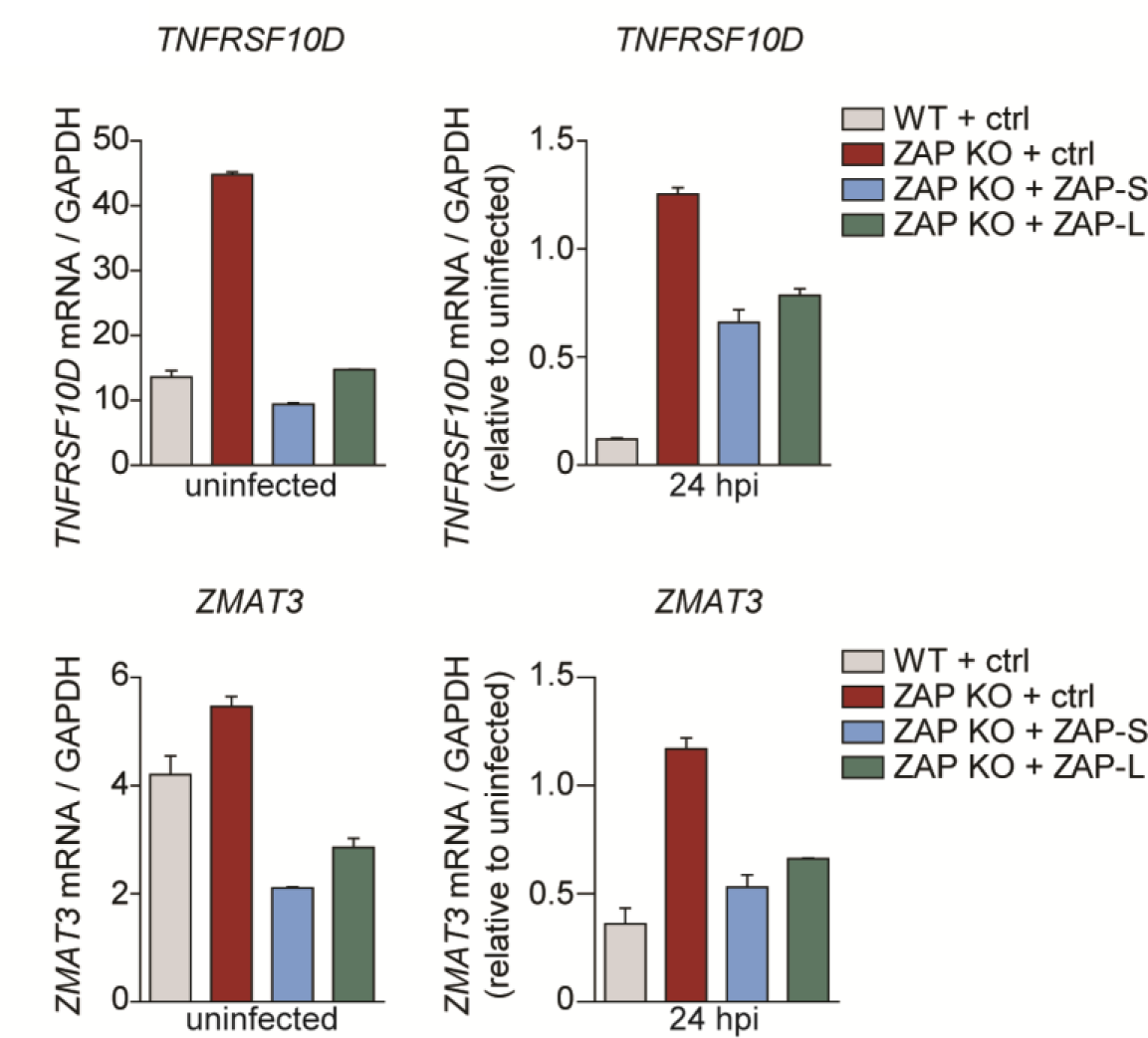
Presence of ZAP-S and ZAP-L leads to reduced *TNFRSF10D* and *ZMAT3* cellular transcripts levels. WT, ZAP KO, and ZAP KO HFF-1 cells expressing either ZAP-S (blue) or ZAP-L (green) were mock treated or infected by centrifugal enhancement with HCMV (MOI 0.1). At 24 hpi, total RNA was extracted and qRT-PCR for *TNFRSF10D* and *ZMAT3* mRNA was performed. Cellular mRNA expression normalised to *GAPDH* is displayed as bar plots showing mean ± S.D. of experimental duplicates. One representative of two independent experiments is shown.

**Figure EV3.**
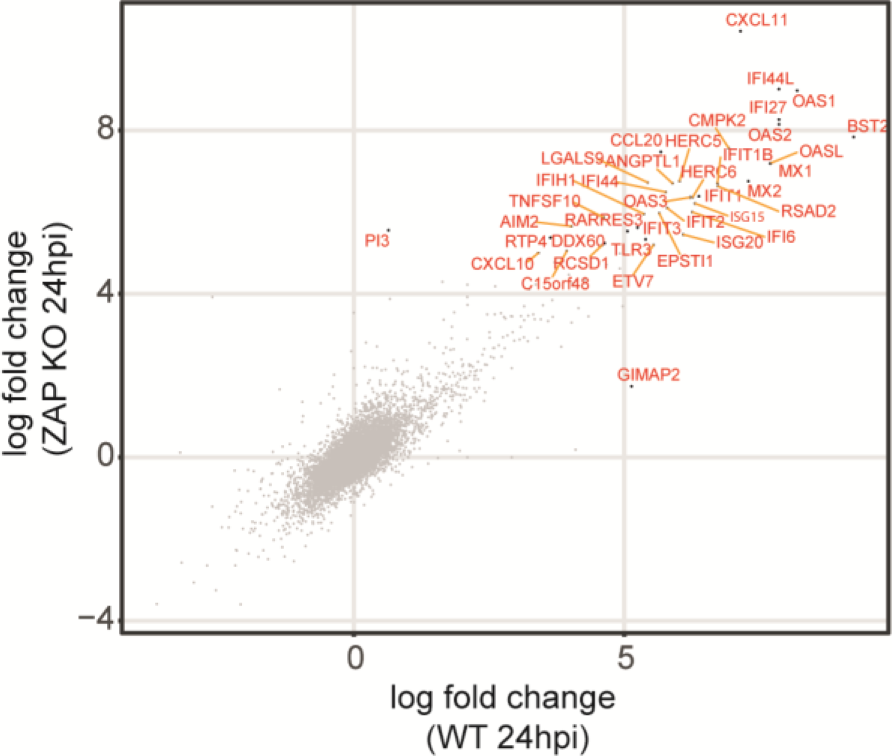
WT and ZAP KO cells show similar induction of ISGs during HCMV infection. WT and ZAP KO HFF-1 cells were untreated or infected by centrifugal enhancement with HCMV (MOI 0.1). Total RNA was extracted at 24 hpi, and lysates were subjected to total transcriptome analysis. Represented are Log_2_ transformed fold changes at 24 hpi compared to untreated cells of WT and ZAP KO (g3) cell lines, calculated using edgeR, and plotted against each other. ISGs are depicted in red.

**Figure EV4.**
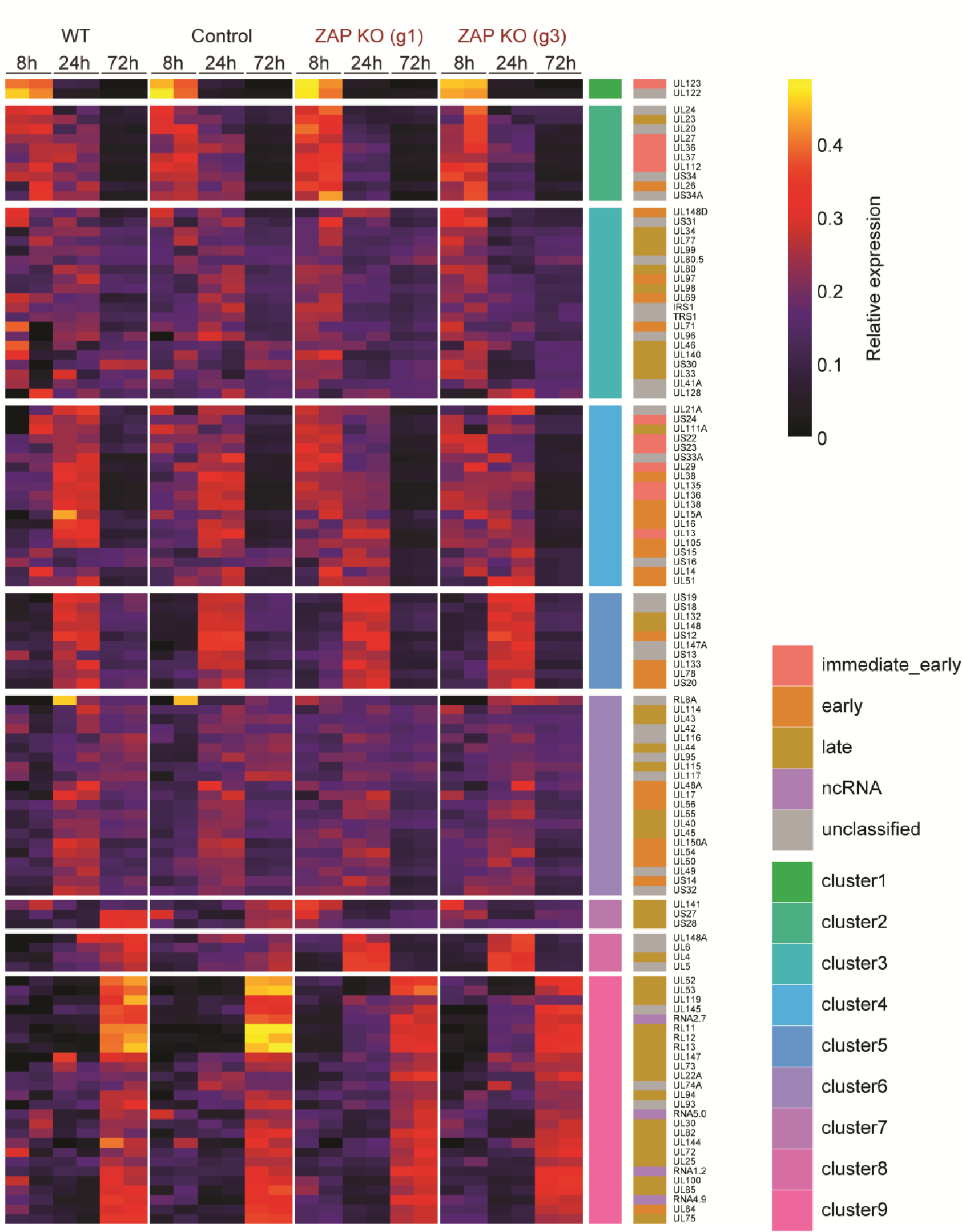
Relative temporal expression levels of HCMV genes above a reasonable expression threshold at 8, 24 and 72 hpi. WT, control, and two independent ZAP KO HFF-1 cell lines were untreated, mock treated, or infected by centrifugal enhancement with HCMV (MOI 0.1). Total RNA was extracted at 8, 24, and 72 hpi, and lysates were subjected to total transcriptome analysis. Expression of HCMV genes was quantified from the RNA-sequencing analysis and relative temporal expression levels calculated by dividing per-sample normalised expression values (fpkm) to the sum of these values from the same gene over six samples of the same cell line. Based on these values, genes were grouped using unsupervised clustering, and the clusters, representing kinetic classes, ordered from immediate early (top) to late (bottom). In addition, shown to the right is the kinetic classification from Weekes *et al*. (2014) (immediate early, early, late) where available, or if the gene codes for a non-coding RNA.

